# Phosphoproteomics identifies the DYRK1B protein kinase as a regulator of processing bodies

**DOI:** 10.64898/2026.04.29.721300

**Authors:** Anne Ashford, Suzan Ber, Miriam S Ems, Emma Duncan, Kathryn Balmanno, Hannah Reeves, Rachael Huntly, Megan A Cassidy, Harvey E Johnston, David Oxley, Thaddeus Mutugi Nthiga, Terje Johansen, Marie Kluge, Ralf Jacob, Matthias Lauth, Simon J Cook

## Abstract

Dual-specificity tyrosine-phosphorylation-regulated kinase 1B (DYRK1B) modulates the cell cycle, cell fate during development, and is deregulated in cancer and metabolic syndrome. However, only a few DYRK1B substrates have been defined, so we undertook a phosphoproteomics screen in cells that exhibit inducible DYRK1B expression. Motif analysis revealed enrichment for proline-directed serine or threonine phosphorylation sites (pSer/pThr-Pro), consistent with the consensus motif of class I DYRKs. Gene ontology analysis revealed enrichment of proteins involved in mRNA binding, mRNA processing and ribonucleoprotein complexes. Several processing body (PB) components, including DCP1A, PATL1(PAT1B), EDC3 and 4E-T, were identified as DYRK1B-inducible phosphoproteins. DYRK1B also co-immunoprecipitated with DCP1A, PAT1B, EDC3, EDC4, DDX6 and XRN1. Super-resolution microscopy demonstrated that DYRK1B co-localised with DCP1A, DCP1B and DDX6 in PBs. Activation of DYRK1B increased PB abundance, whereas inhibition, depletion or knockout of DYRK1B reduced phosphorylation of DCP1A and 4E-T and decreased PB number. Re-expression of wild type, but not kinase-dead, DYRK1B restored PB numbers in knockout cells. These findings reveal novel DYRK1B targets and establish DYRK1B as a regulator of processing body abundance.

**Highlights:**

- DYRK1B induces phosphorylation of a cluster of RNA binding and processing body associated proteins.
- DYRK1B localises to PBs and associates with multiple PB components.
- DYRK1B controls P-body abundance in a kinase-dependent manner.

## INTRODUCTION

The dual-specificity tyrosine-phosphorylation-regulated kinases (DYRKs) are found within the CMGC (CDK, MAPK, GSK and CLK) arm of the eukaryote kinome and are highly conserved throughout evolution (Becker *et al*, 1998; Becker & Joost, 1998; Aranda *et al*, 2011; Han *et al*, 2012). DYRKs require phosphorylation of the second tyrosine in the activation loop tyrosine-X-tyrosine (Y-X-Y) motif to become mature, active kinases (Himpel *et al*, 2001; Lochhead *et al*, 2003). In contrast to most other protein kinases, this activating phosphorylation is not catalysed *in trans* by an upstream ‘activating kinase’ but occurs during translation when DYRKs undergo intramolecular *cis*-autophosphorylation on tyrosine (Lochhead *et al*, 2005; Himpel *et al*, 2000). As a consequence, DYRKs are expressed in their active form, suggesting that regulation of their expression, localisation and/or protein-protein interactions are key for DYRK functions. DYRKs phosphorylate their substrates on serine or threonine residues, with a strong preference for a proline-directed context (pS-P or pT-P) (Soundararajan *et al*, 2013; Himpel *et al*, 2000) like the mitogen-activated protein kinases (MAPKs) and cyclin-dependent kinases (CDKs).

The mammalian DYRKs are divided into class I (DYRK1A and 1B) and class II (DYRK2, 3 and 4) and have roles in transcription, mRNA splicing, cell cycle progression, survival and differentiation (Becker & Joost, 1998; Aranda *et al*, 2011). Both DYRK1A and DYRK1B can drive cell cycle arrest by phosphorylating and degrading cyclin D1 (CCND1) and increasing the abundance of the cyclin-dependent kinase inhibitors p21^CIP1^ and p27^KIP1^ (Ewton *et al*, 2003; Chen *et al*, 2013; Ashford *et al*, 2014; Soppa *et al*, 2014). Much of our knowledge of DYRKs has stemmed from study of *DYRK1A,* which is situated in the Down syndrome (DS) critical region of chromosome 21, is overexpressed in foetal and adult brains of DS individuals (Dowjat *et al*, 2007) and contributes to the clinical features of Down syndrome (Smith *et al*, 1997; Guimera *et al*, 1999; Altafaj *et al*, 2001). Indeed, triplication of *DYRK1A* decreases nuclear CCND1 levels and drives cortical neurogenic defects in a mouse model of DS (Najas *et al*, 2015). Conversely, loss of a single copy of *DYRK1A* in mice leads to increased apoptosis and decreased brain size (Fotaki *et al*, 2002) emphasizing the importance of *DYRK1A* gene dosage and activity. DYRK1B is implicated in several models of differentiation including myogenesis (Deng *et al*, 2003) and adipogenesis (Leder *et al*, 2003). Mutations in *DYRK1B* have been reported in an inherited form of metabolic syndrome associated with early onset coronary artery disease, obesity, hypertension and diabetes (Keramati *et al*, 2014). In addition, *DYRK1B* is amplified (Kuuselo *et al*, 2007; Davis *et al*, 2013) and mutated (Greenman *et al*, 2007) in certain cancers and can promote cell survival (Gao *et al*, 2009; Deng *et al*, 2009) and tumour immune evasion (Ems *et al*, 2025; Brichkina *et al*, 2024)

The preceding studies indicate that the DYRKs control critical cell fate decisions and are deregulated in disease, much like other CMGC kinases. Despite this, relatively few DYRK substrates have been reported that might account for their biological effects, whereas hundreds of substrates are known for ERK1/2 and the CDKs (Yoon & Seger, 2006; Courcelles *et al*, 2013; Malumbres, 2014; Carlson *et al*, 2011; Petrone *et al*, 2016). Most substrates have been reported for DYRK1A and include transcription factors (e.g., NFAT, GLI1), splicing factors (cyclin L2, SF2, SF3), a translation factor (eIF2Bε) and synaptic proteins (dynamin I, synaptojanin I) (Park *et al*, 2009). DYRK1A also phosphorylates Tau (Kimura *et al*, 2007) and this may be relevant to early onset Alzheimer’s disease in DS and a wider role for DYRK1A in neurodegeneration (Park *et al*, 2009). Some substrates are shared by different DYRKs; for example both DYRK1A and DYRK1B phosphorylate CCND1 at T286 to target it for proteasomal degradation (Chen *et al*, 2013; Ashford *et al*, 2014; Soppa *et al*, 2014). Far fewer substrates have been defined for the class II DYRKs, although DYRK2 can phosphorylate p53 at S46 (Taira *et al*, 2010). Beyond DYRK1A, other DYRK family members are increasingly recognized for their roles in neurodevelopmental diseases and cancer, making DYRK kinases potentially attractive therapeutic targets. Identifying new DYRK substrates may provide a molecular basis for DYRK biology but also support drug discovery efforts, since validated substrates may serve as biomarkers of DYRK inhibition.

In addition to their established roles in cell cycle regulation, differentiation and metabolism, DYRK family kinases have been increasingly implicated in the control of membraneless organelles (Álvarez *et al*, 2003; Wippich *et al*, 2013). Membraneless organelles such as processing bodies (PBs), involved in translational repression and RNA turnover (Parker & Sheth, 2007), are dynamic ribonucleoprotein assemblies formed through multivalent interactions between proteins and RNA and are regulated by post translational modifications, including phosphorylation (Rzeczkowski *et al*, 2011; Aizer *et al*, 2013; Chiang *et al*, 2013).

In this study we identify DYRK1B as a PB-associated kinase that directly phosphorylates the PB components DCP1A and 4E-T, promotes phosphorylation of additional PB-associated proteins, and localises to PBs. We further show that DYRK1B kinase activity is required to maintain PB abundance in multiple cell lines, including pancreatic cancer cells. Together these findings establish DYRK1B as a novel regulator of PB abundance.

## RESULTS

### Identification of DYRK1B-induced phosphoproteins by phospho-SILAC mass spectrometry

The DYRKs undergo *cis*-autophosphorylation at a conserved tyrosine in their activation loop during translation so they are active once they are expressed. Consequently, we employed HEK293 cells with tetracycline-inducible expression of DYRK1B (HD1B cells) (Ashford *et al*, 2014) to identify DYRK1B-inducible phosphoproteins using stable isotope labelling using amino acids in cell culture (SILAC). HD1B cells were grown in ^12^C (R0K0) or ^13^C (R6K6) SILAC media for 11 days by which time ^13^C incorporation in R6K6 was 98-99%; R6K6 cells were then treated with tetracycline for a further 8 hours prior to lysis (Fig. 1A). This timepoint was chosen as the earliest point at which DYRK1B expression was maximal and was a trade-off; earlier timepoints would more likely identify proximal DYRK1B targets (e.g., direct DYRK1B substrates) whereas longer timepoints would more likely identify distal phosphorylation events (e.g., those arising from activation of other DYRK1B-dependent protein kinases). An aliquot of each dish was set aside for western blot analysis and confirmed expression of DYRK1B, turnover of CCND1 and phosphorylation of p27^KIP1^ (Fig. 1B) (Ashford *et al*, 2014). The remaining lysates were normalised for protein content, pooled as -Tet/+Tet pairs, reduced and digested with trypsin. Phosphopeptides were enriched using titanium dioxide beads and analysed by LC/MS-MS (Fig. 1A). The full results of this are shown in Table S1. An example trace for a differentially phosphorylated peptide is shown in Fig. EV1; this peptide was identified as being derived from DCP1A, a regulatory subunit that binds to the catalytic subunit DCP2 to form a mRNA decapping complex (Mugridge *et al*, 2016)

**Figure 1:**
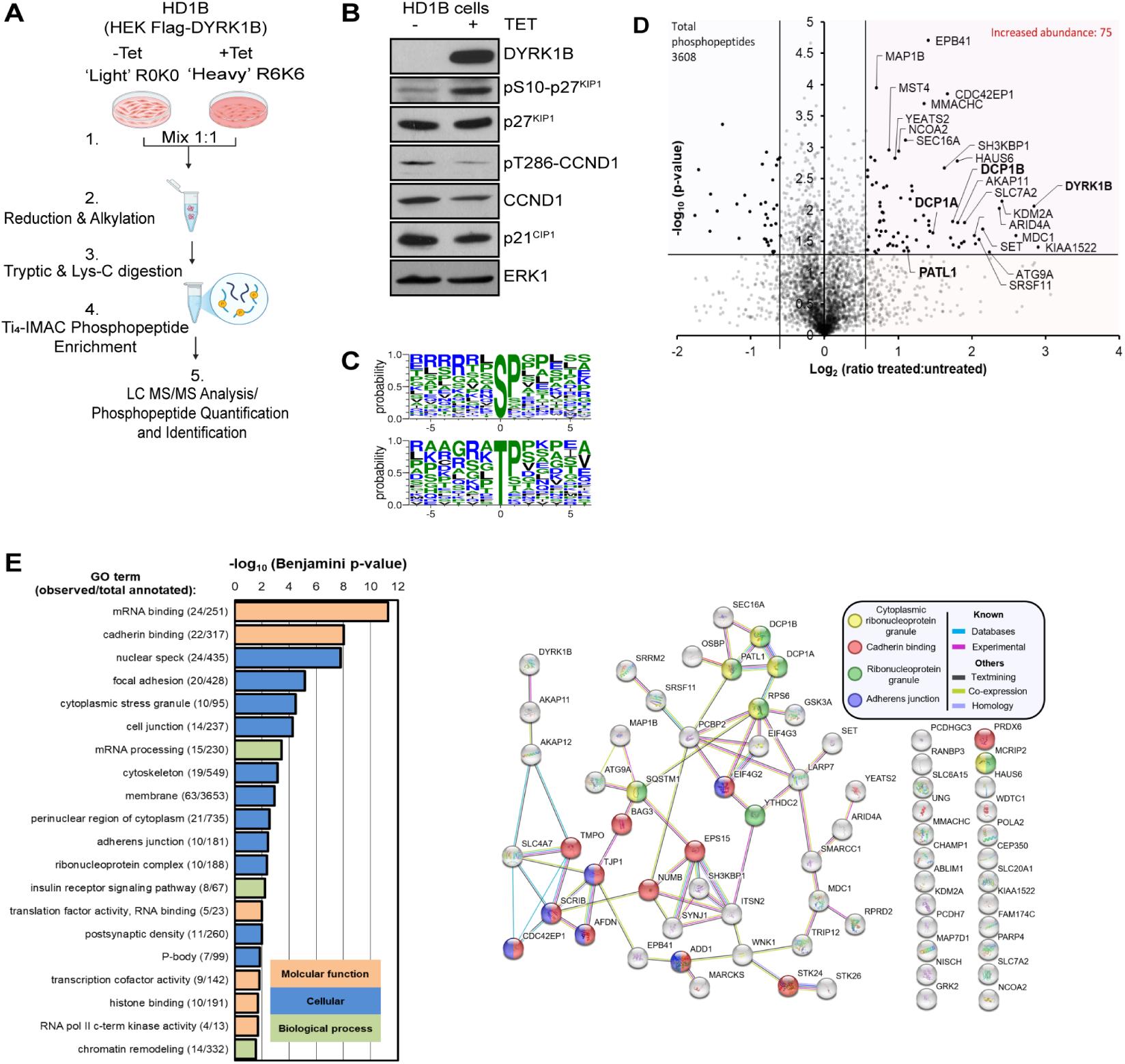
Identification of DYRK1B-inducible phosphoproteins. (A) Experimental workflow for phospho-SILAC analysis of HD1B cells which exhibit Tetracycline inducible DYRK1B expression. Equal amounts of lysate from untreated light labelled cells and Tetracycline-treated heavy labelled cells were combined, digested, phosphopeptides enriched, and analysed by LC-MS/MS. (B) Immunoblot validation of DYRK1B induction after Tetracycline treatment (1 µM, 8 h), with increased p27 phosphorylation and reduced Cyclin D1 levels. (C) Motif analysis of phosphosites increased following DYRK1B induction. (D) Volcano plot detailing SILAC-quantified phosphopeptide abundance changes upon treatment. Phosphopeptides with a fold change >1.5 and *p* < 0.05 are highlighted in red. (E) Functional characterisation of proteins with increased phosphorylation (>1.5-fold, *p* < 0.05). Gene ontology (GO) enrichment analysis (left) identifies significantly overrepresented biological processes. STRING network analysis of the same proteins (right) reveals interaction networks and functional clustering of DYRK1B regulated proteins.

All differentially phosphorylated peptides identified were phosphorylated on Serine or Threonine except one; a peptide derived from DYRK1B itself showed increased phosphorylation of Tyrosine in the activation loop (I**<u>Y</u>**Q**<u>Y</u>**IQSR) (Table S1) consistent with inducible DYRK1B expression. Motif analysis derived from all differentially phosphorylated peptides revealed a strong enrichment for proline-directed sites (Ser-Pro or Thr-Pro) with an additional preference for arginine upstream of the phosphoacceptor site (Fig. 1C). This agrees with previous reports of class I DYRK substrate selectivity (Himpel *et al*, 2000; Campbell & Proud, 2002) suggesting that some of the proteins identified could be direct DYRK1B substrates.

The DYRK1B-inducible phosphoproteins that we identified (Fig. 1D) are involved in a range of biochemical processes including: cell signalling (ADRBK1, AKAP11, GSK3A, NISCH, SCRIB, STK24/MST3, STK26/MST4, SYNJ1, WNK1); protein trafficking (EPS15, SH3KBP1); autophagy and proteostasis (ATG9A, BAG3, SQSTM1); protein ubiquitylation (TRIP12, WDTC1); protein synthesis (RPS6, EIF4G2, EIF4G3); the actin cytoskeleton (ABLIM1, CDC42EP1); cell-cell adhesion (Afadin); microtubule dynamics (KATNA1); metabolite or amino acid transport (SLC4A7, SLC6A15, SLC7A2, SLC20A1); DNA damage checkpoint (MDC1), histone modifications and chromatin remodelling (KDM2A, SET, SMARCC1); regulation of transcription (ARID4A) and cell cycle (MPLKIP, RBL1). Some of these proteins have been reported to be phosphorylated upon DYRK1B overexpression (Dong *et al*, 2020), reinforcing the robustness of our observations. Notably, gene ontology (GO) analysis of DYRK1B-inducible phosphoproteins suggested roles in mRNA binding, mRNA processing and RNP complexes as well as roles in focal adhesions, cadherin binding and adherent junctions (Fig. 1E).

### Validation of Processing body proteins as DYRK1B targets

Prompted by the GO analysis we noted that DCP1A, DCP1B, 4E-T, PAT1B (also known as PATL1) and EDC3 were phosphorylated upon DYRK1B activation (Table S1). These proteins have roles in mRNA decapping (DCP1A/B, EDC3, PAT1B) or translational repression (4E-T) and localise together in cytosolic membraneless organelles or condensates called processing bodies (PBs). Notably, many of the DYRK1B-driven phosphorylation sites on DCP1A, DCP1B, 4E-T and PAT1B identified in the screen were proline-directed (Table S1) agreeing with the DYRK phosphorylation site consensus.

In follow-up experiments, inducible DYRK1B expression led to a time-dependent reduction in electrophoretic mobility of DCP1A and 4E-T in HD1B cells. This shift is typical of phosphorylation and correlated well with the onset of DYRK1B expression (Fig. 2A). The DYRK1B induced mobility shift of DCP1A and 4-ET was reversed by the DYRK1B-selective inhibitor AZ191 (Ashford *et al*, 2014), whereas a catalytically inactive or kinase-dead DYRK1B D239A mutant failed to induce DCP1A or 4E-T mobility (Fig. 2B). Together these results indicate that the DCP1A and 4E-T gel shifts reflect DYRK1B-driven phosphorylation in cells. On conventional SDS-PAGE gels there were very subtle changes for PAT1B and EDC3 whereas Phos-Tag gels revealed clear DYRK1B-dependent phosphorylation of PAT1B and EDC3 and enhanced the phosphorylation-dependent gel shift of DCP1A and 4E-T (Fig. 2C). DYRK1B-dependent phosphorylation of DCP1A was also observed in HeLa cells expressing DYRK1B from the Trex system (HeLa Flp-In T-REx) causing a DCP1A mobility shift and increased phosphorylation of the autophagy cargo receptor p62/SQSTM1 at T269/S272, confirming the p-SILAC data (Fig. 2D). In both cases these phosphorylation events were not observed upon expression of either of two kinase dead mutants, DYRK1B K140R or DYRK1B D239A. DYRK1B expression increased the abundance of DCAF7, a scaffold protein that can recruit class I DYRKs to some of their substrates (Yu *et al*, 2019; Glenewinkel *et al*, 2016). This increase in DCAF7 was also observed with DYRK1B K140R or DYRK1B D239A suggesting that DCAF7 may be stabilized by its interaction with DYRK1B, independent of DYRK1B kinase activity.

**Figure 2:**
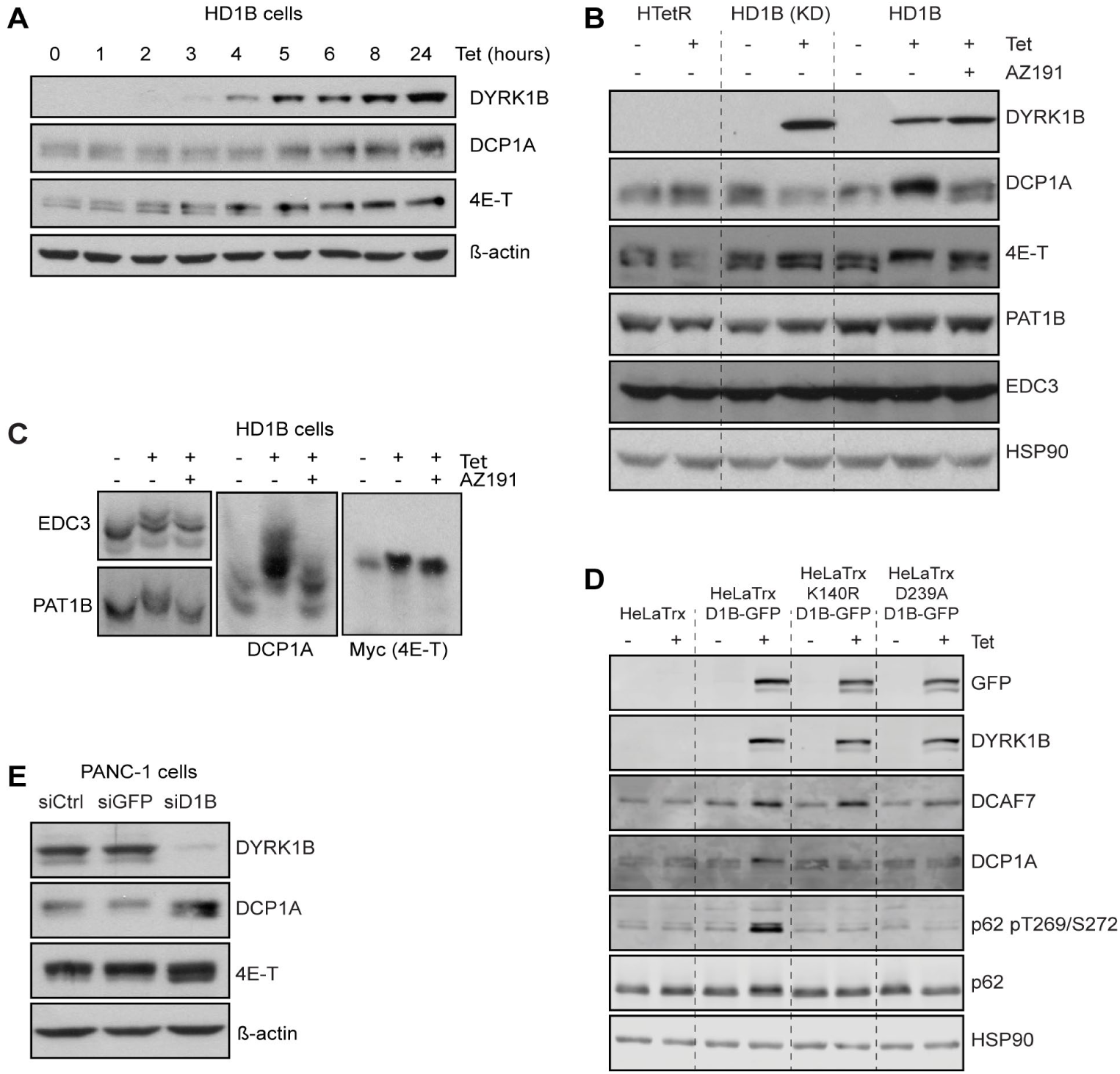
Validation of Phospho-SILAC targets. (A) Time course induction of FLAG-DYRK1B in HD1B cells using Tetracycline (1µM) for up to 24h shows increased phosphorylation of DCP1A and 4E-T targets. Phosphorylation coincides with the expression of DYRK1B protein. (B) Expression of WT but not kinase dead (KD) DYRK1B, upon Tetracycline (1µM) treatment for 24h, phosphorylates DCP1A, 4E-T and PAT1B in HD1B cells. This phosphorylation is inhibited if cells are treated with the DYRK1B inhibitor AZ191 (1µM). (C) Phos-tag gel showing the shift of phosphorylated 4E-T, DCP1A, PAT1B, EDC3 proteins upon DYRK1B expression and the impaired phosphorylation upon treatment with AZ191 (1µM). (D) Expression of WT but not kinase dead (K140R and D239A mutants) EGFP-DYRK1B in HeLa Flp-In T-Rex cells increases phosphorylation of DCP1A and p62 (T269/S272). (E) Knockdown of DYRK1B in PANC-1 cells using siRNA reduces DCP1A and 4E-T phosphorylation.

We have demonstrated that DCP1A, PAT1B, EDC3 and 4E-T underwent phosphorylation upon activation of DYRK1B in cells. To see if DYRK1B was required for their phosphorylation we analysed pancreatic cancer cell lines since *DYRK1B* amplifications are associated with ∼7% of pancreatic cancer (Fig. EV2A). PANC-1 cells harbor a 19q13.1 amplification that includes the *DYRK1B* gene and exhibit greatly elevated expression of DYRK1B relative to BxPC3 cells which lack the 19q13.1 amplification (Fig. EV2B). When PANC-1 cells were transfected with siRNA to knock-down DYRK1B expression, DCP1A and 4E-T mobility on SDS-PAGE increased, consistent with their dephosphorylation (Fig. 2E). Thus, PANC-1 cells were dependent upon endogenous DYRK1B for phosphorylation of these PB proteins.

### DCP1A and 4E-T are direct substrates of DYRK1B

Phos-Tag gels indicated DCP1A underwent multi-site phosphorylation following DYRK1B activation in cells. To assess if DCP1A and 4E-T were direct substrates of DYRK1B we conducted *in vitro* kinase assays with purified components. Purified, recombinant DYRK1B catalysed phosphorylation of both DCP1A and 4E-T *in vitro* and this was inhibited by the inclusion of AZ191 and Harmine, a DYRK1A inhibitor with some activity against DYRK1B (Göckler *et al*, 2009) (Fig. 3A). Phos-Tag analysis of these reactions revealed multiple phosphorylated species of DCP1A, consistent with multi-site modification (Fig. 3B).

**Figure 3:**
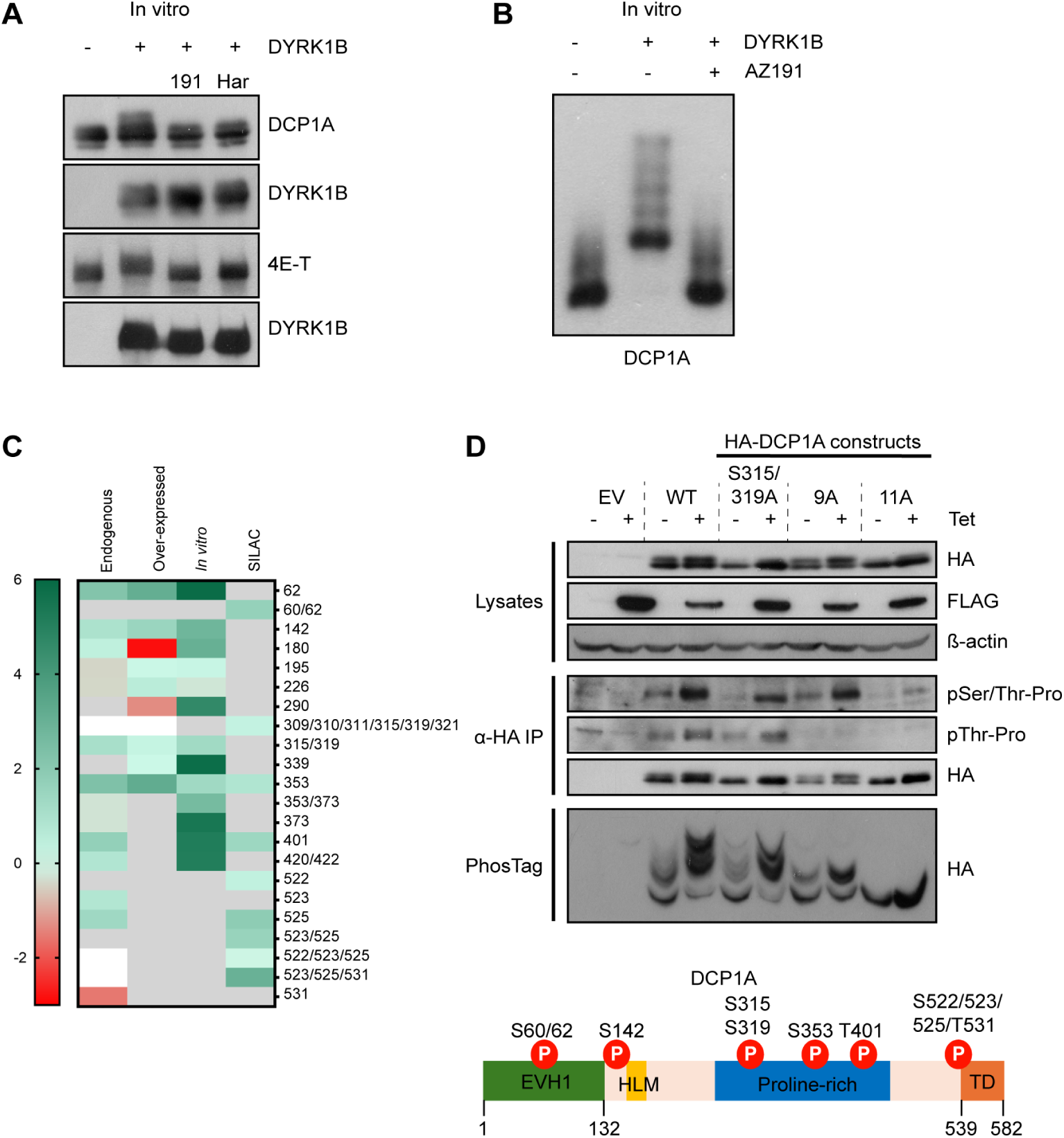
DCP1A and 4E-T are direct substrates of DYRK1B. (A) In vitro kinase assay showing phosphorylation of DCP1A and 4E-T upon addition of recombinant DYRK1B and reduced phosphorylation when co-treated with Class 1 DYRK inhibitors AZ191 (1µM) or Harmine(1µM). (B) Phos-tag gel showing clear shift of *in vitro* phosphorylated DCP1A protein upon DYRK1B addition and the dephosphorylation upon co-treatment with DYRK1B inhibitor AZ191(1µM). (C) Heatmap showing log₂ transformed changes in DCP1A phosphosite abundance upon DYRK1B expression across multiple experimental conditions: DYRK1B-inducible phosphorylation of endogenous DCP1A in HD1B cells; DYRK1B-inducible phosphorylation of overexpressed DCP1A in HD1B cells; direct in vitro phosphorylation by recombinant DYRK1B; and DCP1A phosphorylation sites identified by phospho-SILAC. Values represent log₂ fold changes relative to control for each condition. Grey boxes indicate missing values, corresponding to phosphosites not detected or not quantified in the indicated condition. (D) Pulldown of WT and mutant (S315A/S319A, 9A and 11A phosphonull mutant) HA-DCP1A from HD1B transfected cells, validating multisite phosphorylation sites in DCP1A upon DYRK1B induction.

To define DYRK1B-dependent phosphorylation sites in DCP1A, we used targeted mass spectrometry of DCP1A with Tet inducible DYRK1B in HD1B cells. Fig. 3C shows sites of direct *in vitro* DCP1A phosphorylation by recombinant DYRK1B, phosphorylation sites in over-expressed DCP1A in Tet induced HD1B cells and phosphorylation sites in endogenous DCP1A in Tet induced HD1B cells (also shown in detail in Table S2). Among these DYRK1B-inducible phosphorylation sites S315 and S319 were previously proposed to be phosphorylated by the Pro-directed kinases ERK1/2 (Chiang *et al*, 2013) and JNK (Rzeczkowski *et al*, 2011) and separately to be phosphorylated during mitosis, though the kinase responsible was not defined (Aizer *et al*, 2013).

To assess the contribution of these sites, we generated a S315A/S319A mutant and analysed phosphorylation by mobility shift and phospho-specific antibodies. Mutation of these residues reduced, but did not abolish, the DYRK1B-dependent gel shift and decreased reactivity with the pSer/Thr-Pro antibody, indicating that Ser315 and Ser319 are DYRK1B targets in cells (Fig. 3D). To further evaluate additional sites, we generated a mutant in which nine candidate residues identified by mass spectrometry were substituted with alanine (9A). This mutant showed a marked reduction in DYRK1B-dependent gel shift and loss of pThr-Pro reactivity. Combining these mutations with S315A/S319A (11A mutant) further suppressed phosphorylation, although a residual signal remained, suggesting the presence of additional DYRK1B dependent sites. The residual gel shift and reactivity with the pSer/Thr-Pro antibody suggests that there may be other as yet unidentified DYRK1B phosphorylation sites in DCP1A. These phosphorylation sites are distributed across functionally defined regions of DCP1A, including the N-terminal EVH1 domain (Ser62) which is known to mediate interactions with the DCP2; the central proline-rich region (Ser315/319, Ser353, Thr401 and Ser422); the evolutionarily conserved motif I (Ser142 and Ser180); the C-terminal trimerisation domain (Ser525), indicating that DYRK1B targets multiple structural modules of the protein. These results suggest that Ser62, Ser315/319, Ser353, Thr401 and Ser525 are high confidence DYRK1B phosphorylation sites in cells whilst Ser142, Ser180 and Ser422 are possible DYRK1B phosphorylation sites, all of which are proline-directed.

Collectively, these data identify DCP1A and 4E-T as direct DYRK1B substrates, establish DCP1A as a multi-site target, and show that DYRK1B promotes phosphorylation of multiple PB-associated proteins.

### DYRK1B associates with its substrates and co-localises with them in Processing Bodies

Having identified DCP1A, PAT1B, EDC3 and 4E-T as DYRK1B-inducible phosphoproteins, we next examined whether these proteins physically associate with DYRK1B. FLAG-tagged wild type or kinase-dead (D239A) DYRK1B were transiently expressed and subjected to FLAG immunoprecipitation in both HeLa and HEK293T cells. Across both systems, DYRK1B co-immunoprecipitated with multiple PB components, including endogenous DCP1A, PAT1B and EDC3 in HeLa cells (Fig 4A,B), as well as ectopically expressed HA-DCP1A and EGFP-PAT1B in HEK293T cells using either FLAG or GFP based immunoprecipitation approaches (Fig EV3A,B). In addition, DYRK1B associated with endogenous XRN1, a key 5′–3′ exoribonuclease involved in mRNA decay, in both cell lines (Fig 4B, Fig EV3C) and with the RNA helicase DDX6, which functions in translational repression and RNA turnover, in HeLa cells (Fig 4C).

**Figure 4:**
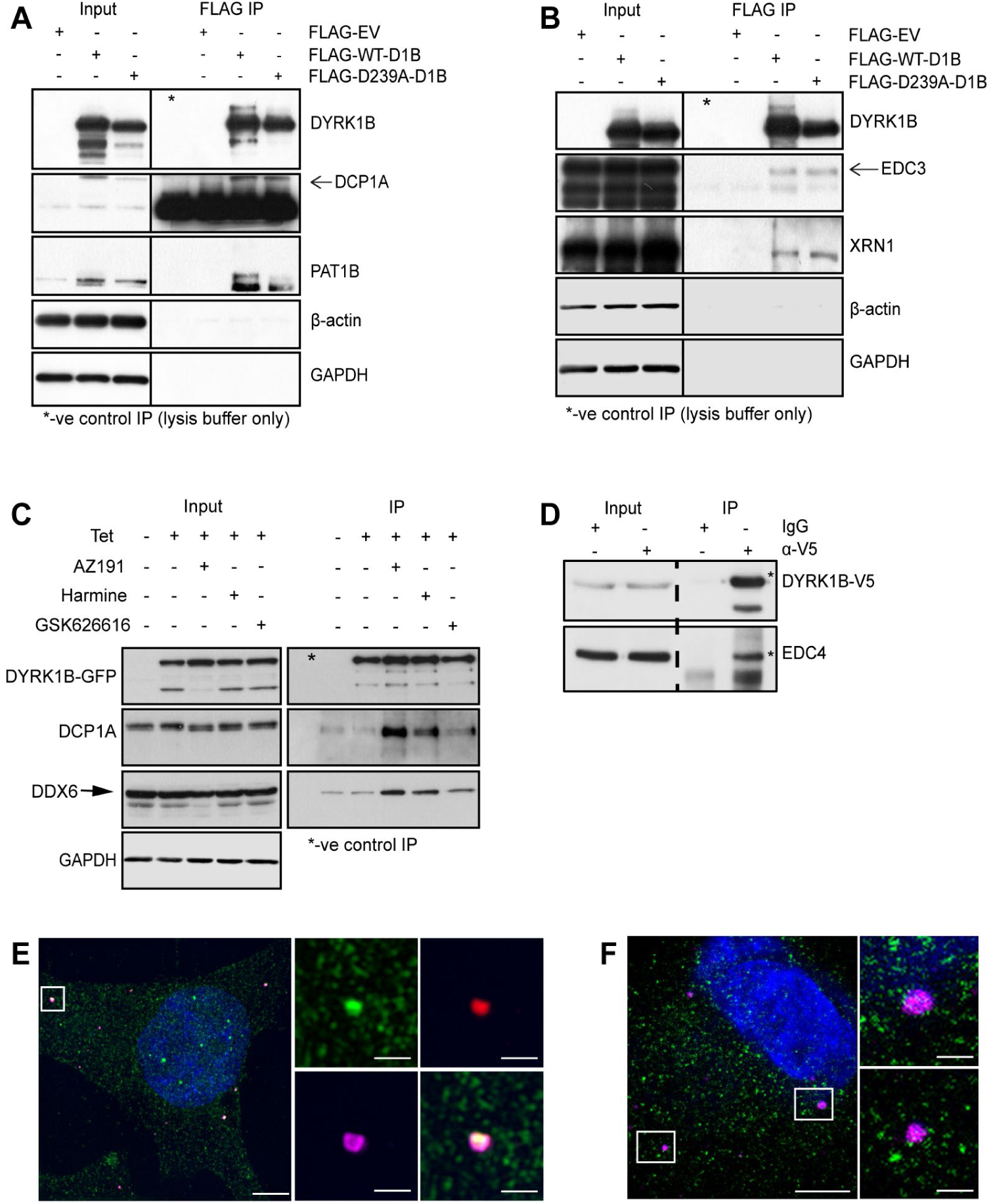
PB components interact with and co-localise with DYRK1B in processing bodies. (A) Immunoprecipitation of FLAG tagged WT or kinase dead DYRK1B (D239A) transfected HeLa cells associates with endogenous DCP1A and PAT1B. (B) Immunoprecipitation of FLAG tagged WT or kinase dead DYRK1B (D239A) transfected HeLa cells associates with endogenous EDC3 and XRN1. (C) Immunoprecipitation of inducibly expressed EGFP-tagged WT DYRK1B from HeLa Flp-In T-Rex cells upon Tetracycline (1µM) induction consecutively treated with DYRK inhibitors for 24h showing interaction with endogenous P-body proteins DCP1A and DDX6. (D) Immunoprecipitation of stably overexpressed V5-tagged DYRK1B in PaTu 8988T cells shows interaction with P-body component EDC4. (E) High-resolution (SIM) image of HeLa Flp-In T-Rex cells expressing EGFP-DYRK1B, showing colocalization with mRNA decapping proteins DCP1A (red) and DDX6 (magenta). (F) Super-resolution (STED) image of endogenous DYRK1B (green) colocalizing with P-body component DCP1B (magenta) in PANC-1 cells. Scale bars represent 5µm, scale bars in zoomed images represent 1µm.

These interactions were retained with the kinase-dead DYRK1B D239A mutant, indicating that binding of DYRK1B to processing body components does not require its catalytic activity (Fig. 4A,B, Fig. EV3A,B,C). To further assess the role of kinase activity in complex formation, HeLa cells expressing EGFP-DYRK1B were treated with DYRK inhibitors prior to immunoprecipitation. Pharmacological inhibition increased the amount of DCP1A and DDX6 co-immunoprecipitating with DYRK1B, with the DYRK1B selective inhibitor AZ191 showing the strongest effect, followed by harmine, whereas the class II DYRK inhibitor GSK626616 had a weaker impact (Fig. 4C). These findings suggest that inhibition of DYRK1B activity stabilises its interaction with PB components. Consistent with these observations, DYRK1B also co-immunoprecipitated with the PB component EDC4 in PaTu 8988T cells expressing V5-tagged DYRK1B (Fig. 4D), supporting a broader association of DYRK1B with the processing body machinery.

To determine whether DYRK1B localizes to PBs, we used super-resolution microscopy to examine its subcellular localisation. This was assessed either by imaging EGFP-tagged DYRK1B in HeLa Flp-In T-REx cells or by staining endogenous DYRK1B in PaTu 8988 cells. PBs were identified using DCP1A and DDX6 co-localised foci in HeLa cells, whereas DCP1B was used as a PB marker in PaTu 8988T cells, all of which are established PB components and label overlapping PB structures, providing comparable measures of PB abundance. DYRK1B co-localised with established PB markers, including DCP1A, DCP1B and DDX6, within cytoplasmic puncta consistent with PB structures (Fig. 4E,F). We also confirmed co-localisation of endogenous DYRK1B with the PB component DCP1B in a second pancreatic cell line, PANC-1 (Fig. EV3D). In addition, live-cell imaging of PaTu 8988T cells expressing DYRK1B-EGFP revealed cytoplasmic GFP puncta that co-localised with mCherry-DDX6 and showed similar dynamic behaviour, consistent with PBs (Movie EV1).

We next compared the localisation of different DYRK family members. EGFP-DYRK1A and EGFP-DYRK1B both showed co-localisation with PB markers DCP1A and DDX6, whereas EGFP-DYRK2 and EGFP-DYRK3 displayed only weak co-localisation (Fig. EV3F). Consistent with this, overexpression of DYRK family members increased phosphorylation of PB proteins, including DCP1A and 4E-T, with the strongest effects observed for DYRK1A and DYRK1B (Fig. EV3E). Taken together, these results show that DYRK1B associates with multiple PB components and localises to PBs.

### DYRK1B regulates Processing body abundance

Considering the data so far we investigated whether DYRK1B might regulate PB abundance. Quantifying PBs (defined as DCP1A and DDX6 positive cytoplasmic foci) in HD1B cells with high content imaging we found that expression of DYRK1B in HD1B cells caused a 3-fold increase in PB abundance that was reversed by AZ191 treatment (Fig. 5A). In contrast, expression of kinase-dead DYRK1B D239A in HD1B cells failed to increase PB abundance (Fig. 5A). These studies were extended to HeLa Flp-In T-REx EGFP-DYRK1B cells; here the basal abundance of PBs was higher and DYRK1B expression caused only a modest increase in PB abundance (Fig. 5B). However, AZ191 treatment decreased both basal and Tet-induced PB abundance suggesting that the higher basal abundance of PBs was in part driven by DYRK1B. These results indicate that DYRK1B expression drives an increase in PB abundance in a kinase-dependent manner (Fig. 5B).

**Figure 5:**
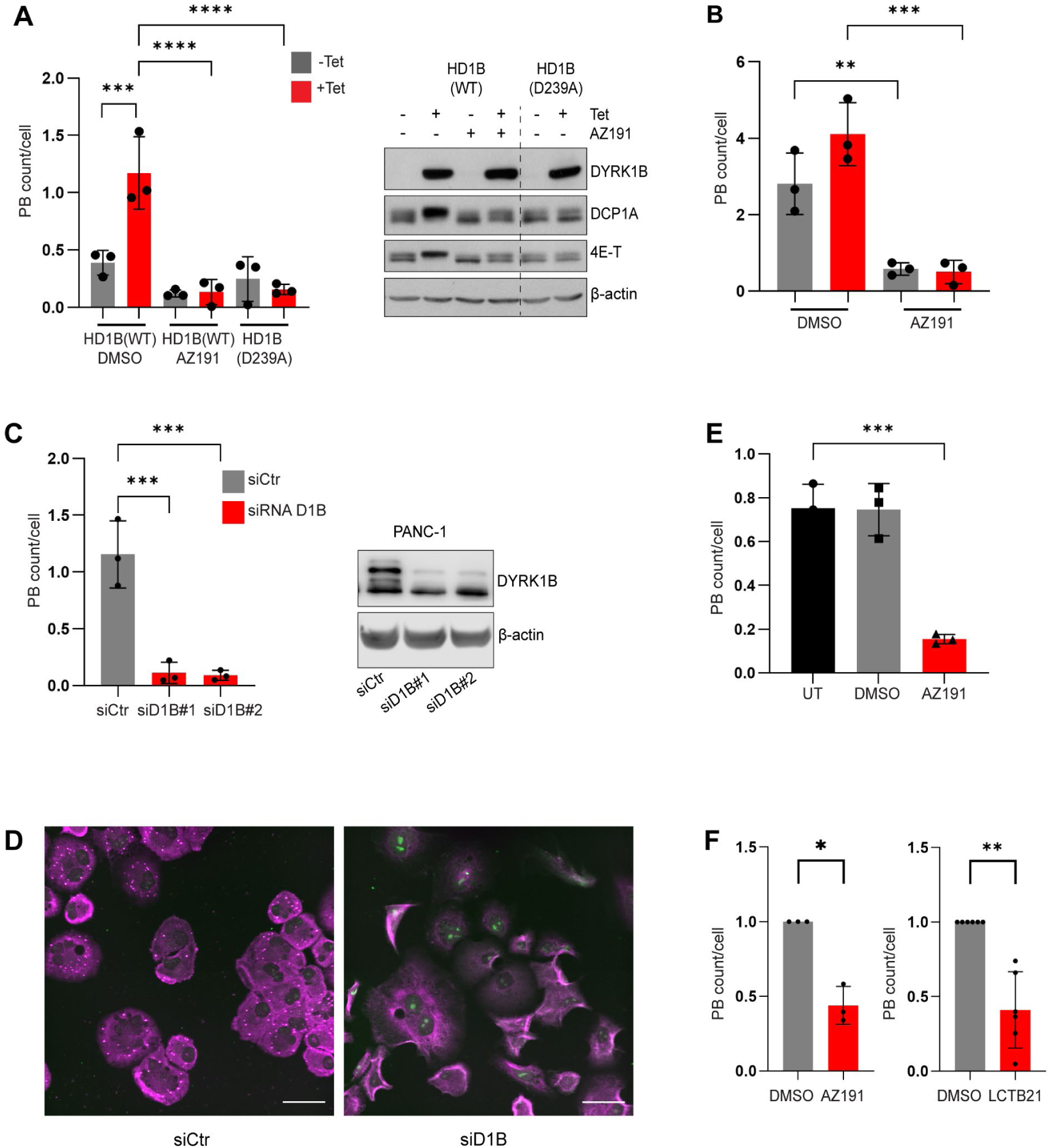
DYRK1B regulates P-body abundance. (A-B) High content quantification of PBs (defined as DCP1A and DDX6 positive foci) in HD1B cells (A) and HeLa Flp-In T-REx EGFP-DYRK1B cells (B). Induction of wild type (WT), but not kinase-dead (D239A) DYRK1B with tetracycline (1 µM, 24 h) increased PB number, which was reduced by AZ191 (1 µM). (C) High-content quantification of PBs (defined as DCP1A and DDX6 positive foci) in PANC-1 cells transfected with two different siRNAs against DYRK1B, showing reduction in P-body number. (D) representative images of high content images of PANC-1 cells used for quantifications in (C), staining of P-body markers DCP1A (magenta) and DDX6 (green). Scale bars, 10µm. (E) High-content quantification of PBs (defined as DCP1A and DDX6 positive foci) in PANC-1 cells treated with DYRK1B inhibitor AZ191(1µM) showing reduction in P-body number. (F) Quantification of PBs using DCP1B as marker showing reduced PB number after 24 h treatment with the DYRK inhibitors AZ191 (1µM) (left, n=3) and LCTB21 (1µM) (right, n=6) in PaTu 8988T cells. Data is presented as mean +/- SD of three independent experiments, unless otherwise stated. p values are calculated using one-way ANOVA (A, B, C, E) with Dunnett’s multiple comparisons test; and unpaired, two-tailed one sample t-test (F) (*p ≤ 0.05, **p ≤ 0.01, ***p ≤ 0.001, ****p ≤ 0.0001).

To address whether DYRK1B was required to maintain PB number in clinically relevant setting we used 2 different pancreatic cancer cell lines, PANC-1 and PaTu 8988T cells. siRNA-mediated knock down of DYRK1B with two different siRNAs in PANC-1 cells, where DYRK1B is amplified (Fig. EV2B), caused a striking >70% reduction in PB abundance as assessed by high content imaging (Fig. 5C,D). This reduction in PB number was phenocopied by the DYRK1B inhibitor AZ191 (Fig. 5E), indicating that this is a kinase-dependent effect of DYRK1B. DYRK1B inhibition in PaTu 8988T cells by either AZ191 or LCTB21 (a DYRK1 inhibitor currently in phase 1 clinical trials) also decreased PB number (Fig. 5F).

Finally, we used CRISPR/Cas9 gene editing to generate PaTu 8988T cells with knockout of DYRK1A or DYRK1B (Fig. 6A). Interestingly, we observed an increase in DYRK1B expression in DYRK1A KO cells, perhaps reflecting a compensatory response and some redundant functions for these two kinases. Despite this, knockout of DYRK1B, but not DYRK1A, significantly reduced the PB number in PaTu 8988T cells (Fig. 6B,E). Re-expression of wild type, but not kinase inactive DYRK1B Y271F/Y273F, restored PB abundance, demonstrating that DYRK1B kinase activity is required for this effect (Fig. 6D,E). These data also further show that DYRK1B, but not the closely related DYRK1A, is required to maintain PB number in pancreatic cancer cells.

**Figure 6:**
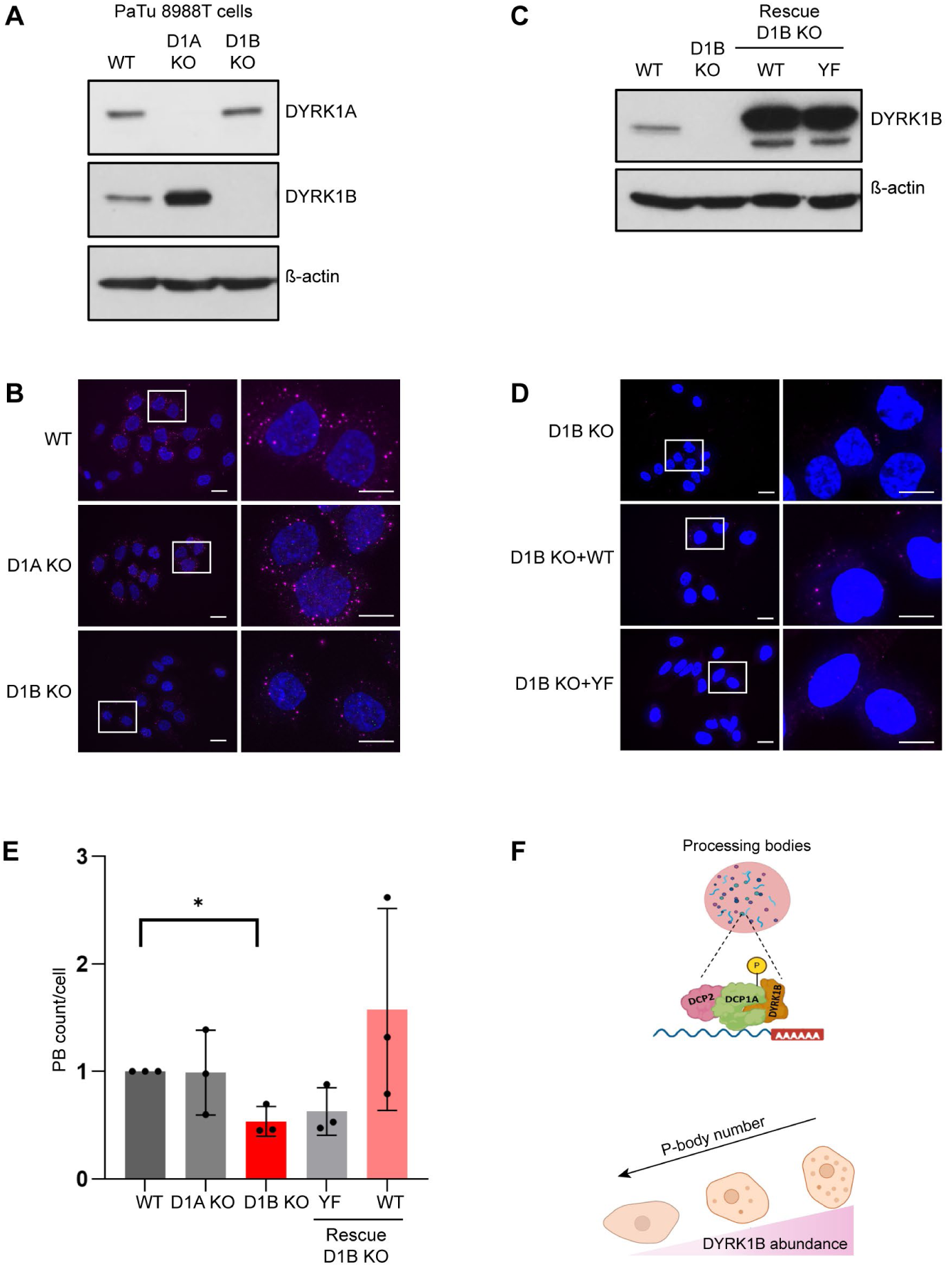
DYRK1B controls P-body abundance in pancreatic cancer cells. (A) Representative western blot images showing loss of DYRK1A (D1A KO) or DYRK1B (D1B KO) in PaTu 8988T pancreatic cancer cells. (B) Representative images of DCP1B foci (magenta) in D1A KO and D1B KO PaTu 8988T cells. The loss of DYRK1B, but not of DYRK1A, leads to reduction in P-body number. Scale bars in the whole cell images represent 20µm and in the zoom panels 10µm. (C) Representative western blot images showing rescue of D1B KO using wild type D1B WT and kinase-dead D1B YF (Y271F/Y273F) DYRK1B in PaTu 8988T cells. (D) Representative images of DCP1B foci (magenta) in D1B KO PaTu 8988Tcells rescued with either wild type D1B WT or kinase-dead D1B YF (Y271F/Y273F). Note the increase of DCP1B foci after rescue with D1B WT but not with D1B YF. Scale bar in the whole cell image represents 20µm and in the zoom panels 10µm. (E) Quantification of DCP1B foci depicted in panels B and D. Data represents mean +/-SD of three independent experiments; p value is calculated using unpaired, two-tailed one sample t-test (*p ≤ 0.05). (F) Model showing previously unknown association of DYRK1B with PBs (created with BioRender). DYRK1B localizes to PBs, associates with several PB components (DCP1A, EDC3, EDC4, PAT1B, XRN1, DDX6) and promotes phosphorylation of these PB-associated proteins. DYRK1B kinase activity is required to maintain PB abundance.

In summary, DYRK1B localises to processing bodies, associates with multiple PB components, including DCP1A, EDC3, EDC4, PAT1B, XRN1 and DDX6, and promotes phosphorylation of DCP1A, EDC3, EDC4, PAT1B and 4E-T. Moreover, DYRK1B kinase activity promotes PB abundance (Fig. 6F).

## DISCUSSION

To identify DYRK1B-inducible phosphoproteins, including DYRK1B substrates, we employed an unbiased phospho-SILAC screen. This analysis identified targets in pathways previously linked to DYRK1B, including cell cycle regulation (MPLKIP, RBL1) and chromatin and transcriptional control (MDC1, KDM2A, SET, ARID4A) (Chen *et al*, 2013; Dong *et al*, 2020, 2021; Ewton *et al*, 2003), while also expanding the DYRK1B phosphoproteome to include proteins involved in translational regulation (RPS6, EIF4G2), autophagy (ATG9A, SQSTM1), transporter function (SLC4A7, SLC6A15, SLC7A2), and cytoskeletal organisation (Afadin, ABLIM1, CDC42EP1, KATNA1). The detection of previously validated substrates (Dong et al, 2020) supports the robustness of our study. A novel finding from this analysis was the enrichment of proteins associated with mRNA processing and ribonucleoprotein (RNP) complexes, particularly core components of processing bodies (PBs), including DCP1A, DCP1B, PAT1B, EDC3 and 4E-T. Further experiments confirmed this connection, demonstrating that DYRK1B activity promotes phosphorylation of these PB-associated proteins in cells and is required to maintain their basal phosphorylation (Fig. 2, EV2). *In vitro* kinase assays further established DCP1A and 4E-T as direct DYRK1B substrates, with DCP1A undergoing multi-site phosphorylation at proline- directed motifs (Fig. 3). Together, these findings identify PB components as a previously unrecognised class of DYRK1B substrates.

We had anticipated that a phosphorylation defective DCP1A mutant might exert an interfering effect when expressed in cells, perhaps by competing with wild type DCP1A to access DCP2, or other regulators. Although mutation of multiple DYRK1B-dependent phosphorylation sites in DCP1A substantially reduced phosphorylation, expression of these mutants did not produce an observable phenotype under the conditions tested. One possible explanation is functional redundancy with DCP1B, which shares significant sequence similarity with DCP1A and contains many of the identified phosphorylation sites. DCP1B also scored in our phospho-SILAC screen and co-localised with DYRK1B in pancreatic cancer cells (Fig. 4E,F). It is therefore plausible that compensation by DCP1B limits the impact of DCP1A phospho mutants, and that combined perturbation of both proteins may be required to reveal functional consequences.

A central finding of this study is that DYRK1B kinase activity promotes and maintains PB abundance. Several DYRK1B substrates localised to PBs, DYRK1B co-immunoprecipitated with multiple PB components (Fig. 4, EV3) and our data support an association of DYRK1B with these structures (Fig. 4). PBs are dynamic ribonucleoprotein assemblies formed through multivalent interactions, with core factors such as EDC4, DDX6, 4E-T (Standart & Weil, 2018) and LSM14 acting as nucleators (Franks & Lykke-Andersen, 2008; Jonas & Izaurralde, 2013; Parker & Sheth, 2007). Disruption of these interactions can impair PB assembly (Brandmann *et al*, 2018), suggesting that modulation of these networks could influence PB formation. Expression of wild type, but not kinase-inactive DYRK1B, increased PB number (Fig. 5), while depletion or knockout of DYRK1B reduced PB abundance in pancreatic cancer cells (Fig. 5,6). This effect was specific to DYRK1B, as DYRK1A did not compensate for its loss. Although DYRK1A phosphorylated PB proteins and localised to PBs when overexpressed, it did not rescue PB abundance in DYRK1B knockout cells, indicating that the two kinases are not functionally redundant in this context. Together, these observations demonstrate that DYRK1B kinase activity is required to maintain PB homeostasis. It is therefore notable that EDC4, DDX6 and 4E-T, which serve as PB nucleators (Franks & Lykke-Andersen, 2008; Jonas & Izaurralde, 2013; Parker & Sheth, 2007), are also DYRK1B substrates or DYRK1B interacting proteins.

The assembly of membraneless organelles such as PBs is driven by multivalent interactions among protein domains, intrinsically disordered regions (IDRs) and RNA (Banani *et al*, 2017, 2016), and is highly sensitive to post-translational modifications (Bah & Forman-Kay, 2016; Hofweber & Dormann, 2018). Phosphorylation can modulate condensate behaviour in context-dependent ways, either promoting or inhibiting assembly. For example, multisite phosphorylation of FUS inhibits condensation (Monahan *et al*, 2017), whereas phosphorylation of FMRP promotes condensate formation (Tsang *et al*, 2019). EDC3 phosphorylation has also been linked to PB regulation in cancer cells (Bearss *et al*, 2021). Interestingly, phosphorylation of DCP1A by JNK or ERK1/2 has been associated with PB dissolution (Yu *et al*, 2024; Rzeczkowski *et al*, 2011; Chiang *et al*, 2013). In this context, DYRK1B dependent phosphorylation of multiple PB components, including DCP1A, 4E-T and PAT1B, may influence their interaction properties and contribute to PB assembly or stability, although the precise mechanism remains to be defined.

Accumulating evidence now supports a wider role for DYRK family kinases as regulators of biomolecular condensate homeostasis (Álvarez *et al*, 2003; Yu *et al*, 2019; Gallo *et al*, 2023). For example, DYRK1A has been shown to localise to and regulate nuclear speckles (Álvarez *et al*, 2003), and DYRK3 regulates stress granule dynamics and may act more broadly as a “dissolvase” of membraneless compartments (Wippich *et al*, 2013). Together with our findings, these studies suggest that DYRK kinases may function as conserved signalling regulators of condensate assembly through multisite phosphorylation of IDR-rich proteins.

Our results identify regulation of PB abundance as a previously unrecognised aspect of DYRK1B biology. While PBs were initially characterised as sites of mRNA decay, they are now understood as dynamic hubs involved in translational control and stress responses (Anderson *et al*, 2015; Bearss *et al*, 2021; Hardy *et al*, 2017). DYRK1B is amplified in a subset of pancreatic cancer and its expression correlates with poor prognosis (Kuuselo *et al*, 2007). Inhibition of DYRK1B has been proposed as a therapeutic strategy to target quiescent tumour cell populations (Deng *et al*, 2009) and modulate immune evasion (Brichkina *et al*, 2024). By regulating the phosphorylation of core PB components, DYRK1B may influence the stability of these condensates and thereby contribute to the adaptive capacity of cancer cells under stress. In this context, phosphorylation of DCP1A and 4E-T may also provide a useful readout or biomarker of DYRK1B activity in cells. In summary, we identify DYRK1B as a non-redundant regulator of PB homeostasis and establish PB components as a novel class of DYRK1B substrates. These findings extend the DYRK1B interactome and suggest a mechanistic link between DYRK1B activity and post-transcriptional regulation of gene expression in cancer.

### Limitations of the study

Although phospho-SILAC identified DYRK1B-inducible phosphorylation events, these likely include both direct substrates and indirect targets. We have however validated a subset of these targets as direct DYRK1B substrates with in vitro assays. Whilst our early experiments employed inducible overexpression of DYRK1B, which may not fully reflect physiological DYRK1B activity, we also studied cancer cell models in which endogenous DYRK1B is amplified (PANC-1 cells) or not (PaTu 8988T) and employed DYRK1B kinase inhibitors, DYRK1B siRNA and CRISPR/Cas9-generated DYRK1A and DYRK1B knockout cells. While we demonstrate that DYRK1B regulates processing body abundance, the mechanistic link between phosphorylation of PB components and condensate assembly remains unresolved. In addition, the functional consequences of DYRK1B activity on mRNA turnover, translation, or transcript specific regulation were not addressed. Finally, all experiments were performed in cell lines, and the in vivo relevance of DYRK1B-dependent PB regulation, particularly in cancer contexts, remains to be defined.

## Supporting information

Live cell imaging PaTu 8988T cells DYRK1B GFP-mCherryDDX6

Supplementary Table S1 DYRK1B SILAC phosphoproteomics

Supplementary Table S2 DCP1A targeted phosphoproteomics

## RESOURCE AVAILABILITY

## Lead contact

Further information and requests for resources should be directed to the lead contact, Simon Cook

## Materials availability

All the materials generated in this study are available upon request to the lead contact.

## Data availability

- All the phospho-SILAC and targeted phosphoproteomics results are available in **Supplemental Table S1 and Table S2.** Raw phospho-SILAC and targeted phosphoproteomics data are available via ProteomeXchange with identifier PXD077447.

## ACKNOWLEDGEMENTS

We would like to thank Nancy Standart (University of Cambridge); Martin Turner, Claudia Ribeiro de Almeida, Ian McGough (Babraham Institute); and members of the Cook Group past and present for encouragement and useful discussions. We are also grateful to excellent staff in the Mass Spectrometry Facility and Imaging Facility at the Babraham Institute, supported by a Core Capability Grant from UKRI-BBSRC. Work in the Cook Group was supported by a collaborative PhD studentship supported by the Biotechnology and Biological Sciences Research Council (BBSRC) and AstraZeneca which was awarded to AA and grants BB/L008793/1, BB/P007015/1, BBS/E/B/000C0417 and BBS/E/B/000C0433 from the UKRI-BBSRC. Work in the Lauth group was funded by a grant from the German Research Foundation (Nr. LA2829/15-1).

## AUTHOR CONTRIBUTIONS

**Anne Ashford:** Conceptualization; Investigation; Data curation; Formal analysis; Validation; Visualization; Writing-review and editing. **Suzan Ber:** Conceptualization; Investigation; Data curation; Formal analysis; Validation; Visualization; Writing-original draft; Writing-review and editing. **Miriam Ems:** Investigation; Data curation; Formal analysis; Validation; Visualization; Writing-original draft. **Emma Duncan:** Investigation; Data curation; Formal analysis; Validation; Visualization. **Kathryn Balmanno:** Investigation; Formal analysis. **Hannah Reeves:** Formal analysis; Data curation. **Rachael Huntly:** Investigation; Formal analysis. **Megan Cassidy:** Investigation; Formal analysis. **Harvey Johnston:** Data curation; Formal analysis. **David Oxley:** Investigation; Data curation; Formal analysis. **Thaddeus Mutugi Nthiga:** Investigation; Formal analysis. **Terje Johansen:** Supervision; Writing-review and editing. **Marie Kluge:** Investigation; Validation; Visualization **Ralf Jacob:** Visualization **Matthias Lauth:** Conceptualization; Formal analysis; Supervision; Funding acquisition; Project administration; Writing-original draft; Writing-review and editing. **Simon J Cook:** Conceptualization; Formal analysis; Supervision; Funding acquisition; Project administration; Writing-original draft; Writing-review and editing.

## STAR METHODS

## KEY RESOURCES TABLE

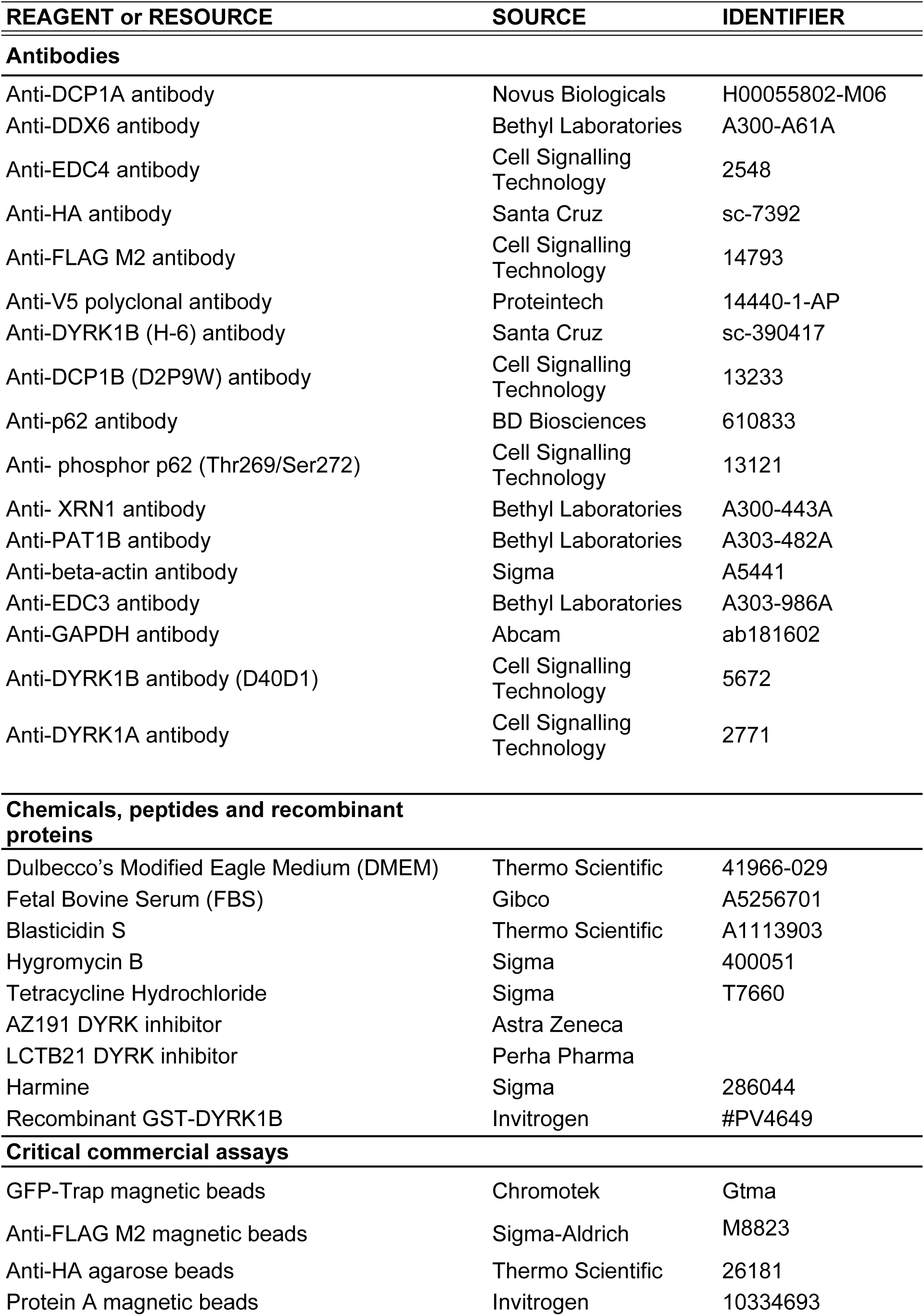
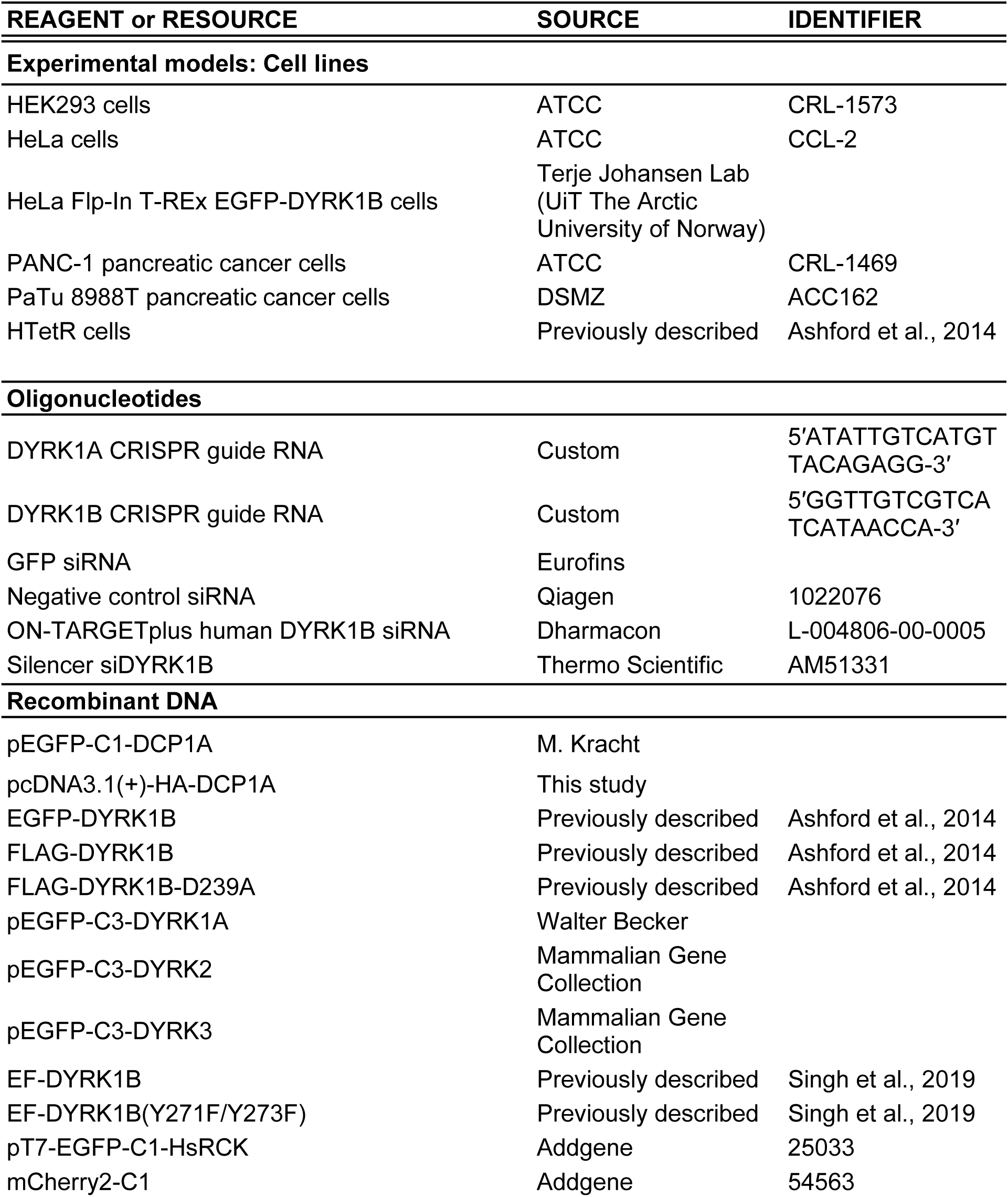
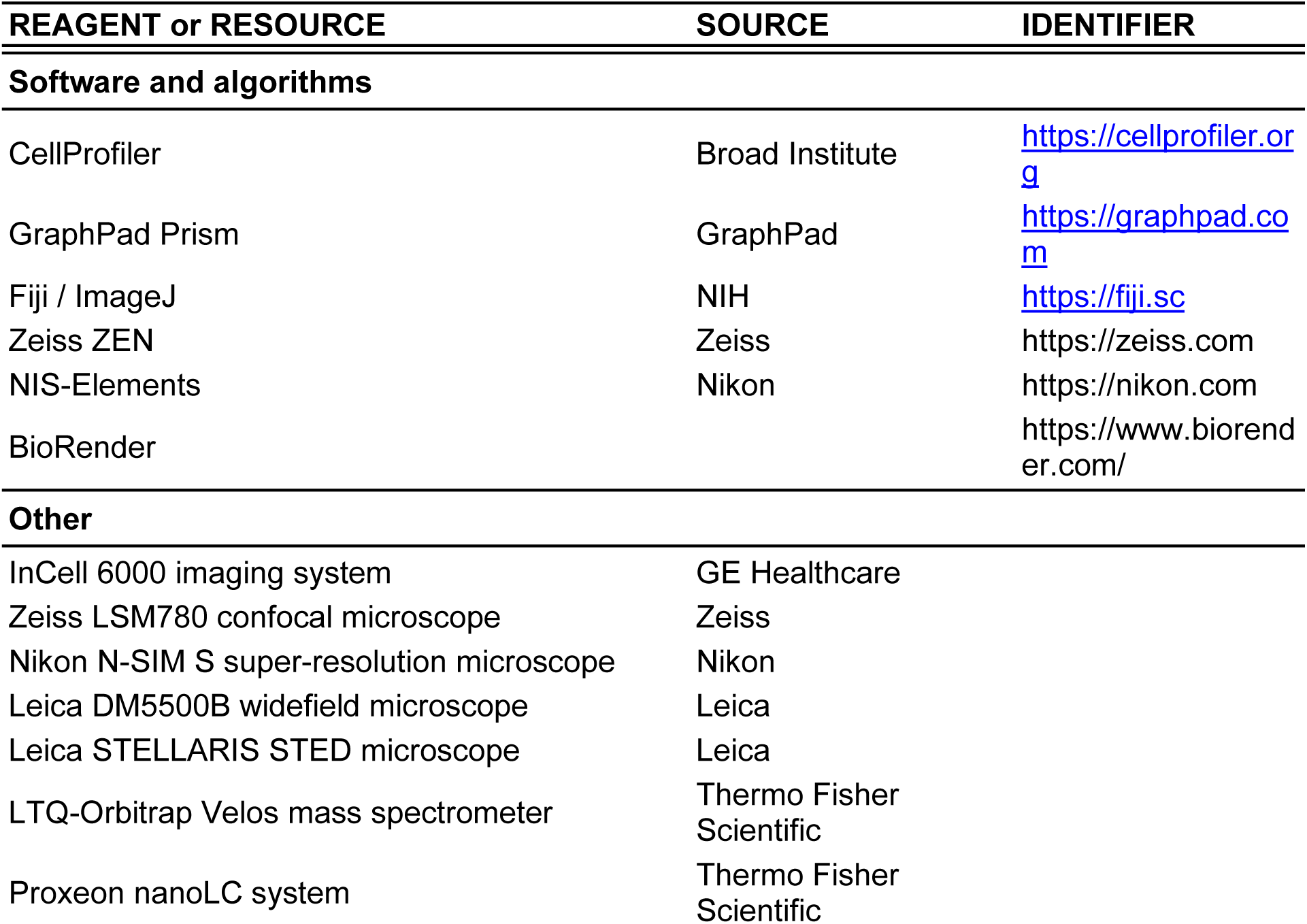

## METHOD DETAILS

### Cell lines and culture conditions

HEK293, PANC-1, HTetR, HD1B and HD1B(KD) cells, including the normal growth media used for the tetracycline-inducible HEK293 derivatives, have been described previously (Ashford *et al*, 2014). DYRK1B expression in HD1B or HD1B(KD) cells was induced by addition of tetracycline to a final concentration of 1 μg/ml.

HeLa cells were obtained from ATCC and maintained in Dulbecco’s modified Eagle’s medium (DMEM) supplemented with penicillin (100 U/ml), streptomycin (100 μg/ml), L-glutamine (2 mM) and fetal bovine serum (FBS; 10% v/v). HeLa Flp-In T-REx cells conditionally expressing EGFP-DYRK1B were maintained in DMEM supplemented with penicillin (100 U/ml), streptomycin (100 μg/ml), FBS (10% v/v) and blasticidin (7.5 μg/ml). The generation of these cells was as follows: DYRK1B cDNA was cloned from pDONR221-DYRK1B (obtained from Harvard plasmids repository) into pDest-Flp-In-EGFP vector by Gateway recombination cloning (Alemu *et al*, 2012). pDest-FlpIn-EGFP-DYRK1B was then co-transfected with recombinase expression plasmid pOG44 into the HeLa Flp-In T-REx cells. After 48 h, cells with the gene of interest integrated into the FRT site were selected with 200 mg/ml of hygromycin and 7.5 mg/ml blasticidin. Hygromycin resistant cells were then expanded in the selection media and later tested for expression by immunoblotting and immunofluorescence. EGFP-DYRK1B expression was induced with tetracycline (1 μg/ml) for the indicated times. PANC-1 and PaTu 8988T pancreatic cancer cells were handled as described previously (Brichkina *et al*, 2024). Where indicated for imaging experiments in pancreatic cancer cells, medium was replaced after 24 h with DMEM containing 0.5% FBS and 1% penicillin/streptomycin, and cells were then left untreated or treated for 24 h with vehicle (0.1% DMSO), AZ191 (1 μM) or LCTB21 (1 μM).

All cell lines were routinely passaged before reaching 80% confluence, maintained at 37°C in a humidified atmosphere containing 5% CO2, and confirmed to be mycoplasma-negative before experiments.

### Generation of knockout and rescue cell lines

CRISPR-mediated depletion of DYRK family kinases in PaTu 8988T cells was performed as described previously (Brichkina *et al*, 2024). The guide sequences used were 5 ′ -ATATTGTCATGTTACAGAGG-3 ′ for human DYRK1A and 5 ′ - GGTTGTCGTCATCATAACCA-3′ for human DYRK1B.

For rescue experiments, DYRK1B-knockout PaTu 8988T cells were transfected with EF-DYRK1B or EF-DYRK1BY271F/Y273F (Singh *et al*, 2019) using Helix-In transfection reagent (Oz Biosciences) for 48h and then selected with blasticidin until single clones were obtained.

### Plasmids, mutagenesis, transfection and siRNA

pEGFP-C1-DCP1A was kindly provided by M. Kracht and was subcloned into pcDNA3.1(+) to generate pcDNA3.1(+)-HA-DCP1A. DCP1A phosphosite mutants were generated by site-directed mutagenesis to produce the S315A/S319A double mutant, the 9A mutant (S60A/ S62A/ S142A/ S353A/ T401A/ S522A/ S523A/ S525A/ T531A) and the 11A mutant (S60A/ S62A/ S142A/ S315A/ S319A/ S353A/ T401A/ S522A/ S523A/ S525A/ T531A).

Full-length EGFP-DYRK1B, FLAG-DYRK1B and FLAG-DYRK1B-D239A constructs have been described previously (Ashford *et al*, 2014). pEGFP-C3-DYRK1A was kindly provided by Walter Becker (Institute of Pharmacology and Toxicology, Aachen University), and DYRK2 and DYRK3 coding sequences from the Mammalian Gene Collection were subcloned into pEGFP-C3. HeLa Flp-In T-REx EGFP-DYRK1B kinase-dead derivatives were generated from the parental EGFP-DYRK1B line by Q5 site-directed mutagenesis (NEB) using the primers listed in Key Resources table.

For transient DNA transfection, cells at approximately 70% confluence were transfected with JetPrime (Polyplus) according to the manufacturer’s instructions. In a 12-well format, 0.5μg DNA was mixed with 75μl JetPrime buffer and 2μl reagent, incubated for 10 min at room temperature, added to cells for 6 h, and then replaced with fresh complete medium for a further 24–48 h.

For knockdown experiments, GFP siRNA (Eurofins), a negative control siRNA (Thermo Fisher Scientific), ON-TARGETplus human DYRK1B siRNA (Dharmacon) and Silencer siDYRK1B (Thermo Fisher Scientific) were used at a final concentration of 30nM. siRNAs were transfected using Lipofectamine RNAiMAX according to the manufacturer’s instructions.

### SILAC phosphoproteomics

HD1B cells were grown in SILAC DMEM (Dundee Cell Products) supplemented with dialysed FBS (10%, 10 kDa cut-off), L-proline (84 mg/L), L-glutamine (2 mM), penicillin, streptomycin, blasticidin and zeocin, and containing either light arginine/lysine (R0K0) or heavy 13C-arginine/13C-lysine (R6K6). Cells were maintained for at least 10 doublings, which gave 98–99% heavy amino-acid incorporation. DYRK1B expression was induced in the heavy-labelled cells by tetracycline (1 μg/ml) for 8h.

Cells were washed briefly in PBS and lysed in ice-cold modified TG lysis buffer (20 mM Tris-HCl, pH 7.5, 137 mM NaCl, 1 mM EGTA, 10 mM EDTA, 1% Triton X-100, 10% glycerol, 1.5 mM MgCl2, 1 mM sodium orthovanadate, 1 mM PMSF, 10 μg/ml leupeptin, 10 μg/ml aprotinin and 50 mM NaF). After clarification (12,000 × g, 10 min, 4°C), equal amounts of protein from control light-labelled and DYRK1B-induced heavy-labelled lysates were combined.

Aliquots of the lysates containing 100µg of protein were precipitated with acetone (4 vols) for 1h at -20°C. The precipitated proteins were solubilised in 25mM ammonium bicarbonate/6M guanidine hydrochloride/10mM DTT (100µL) at 50°C for 1h, then cooled to room temperature and alkylated with iodoacetamide for 30 min in the dark. The S-carbamidomethylated proteins were again precipitated with cold acetone and then solubilised in 25mM ammonium bicarbonate/4M guanidine hydrochloride containing Lys-C protease (1µg). After 1h, the samples were diluted 10-fold with 25mM ammonium bicarbonate containing trypsin (2µg) and incubated for 16h at 50°C. The digestion was terminated by the addition of 10% aqueous trifluoroacetic acid to a final concentration of 0.5%. Phosphopeptides were extracted from the digests with titanium dioxide beads (GL Sciences), eluted from the beads with 5% ammonium hydroxide, then dried and resuspended in 0.1% trifluoroacetic acid.

Phosphopetides were analysed by LC-MS on a Thermo LTQ-Orbitrap Velos mass spectrometer coupled to a Proxeon nanoLC system. Phosphopeptides were separated on a reversed-phase column (0.05 x 500mm Reprosil C18aq 3µm) at a flow rate of 80 nL/min with a gradient of 0-40% acetonitrile (containing 0.1% formic acid) in 8h. The mass spectrometer scan cycle comprised a high resolution (30,000) survey scan followed by up to 20 MS/MS scans (in HCD mode at 7,500 resolution), with 120 seconds dynamic exclusion of former target ions.

The mass spectral data were searched against the human entries in the Uniprot database using Mascot software and the search results processed using Proteome Discoverer to extract SILAC ratios. Differentially regulated phosphopeptides were subsequently used for motif analysis and for downstream enrichment and interaction analyses shown in Fig. 1.

### Immunoblotting, immunoprecipitation and Phos-tag analysis

For standard immunoblotting, cells were lysed in ice-cold TG lysis buffer, clarified by centrifugation and quantified before addition of Laemmli sample buffer. In PaTu 8988T experiments, samples were lysed directly in 1× Laemmli buffer. Proteins were resolved by SDS-PAGE, transferred to Immobilon P/PVDF membranes, blocked in TBS containing 0.1% Tween-20 and 5% milk, and probed with the indicated primary and HRP-conjugated secondary antibodies. Signals were visualised by enhanced chemiluminescence. Where indicated, 50 μM Phos-tag was included in the resolving gel.

For HA-DCP1A pull-downs, HD1B cells expressing HA-tagged wild type or mutant DCP1A were induced with tetracycline (1 μM) for 24h to express FLAG-DYRK1B, lysed in TG buffer, and incubated with anti-HA agarose for 3 h at 4°C. For endogenous EDC4 immunoprecipitation, lysates prepared in IP buffer (50 mM Tris-HCl pH 7.5, 150 mM NaCl, 1 mM EDTA, 0.1% Triton X-100, 10% glycerol plus phosphatase and protease inhibitors) were incubated with anti-EDC4 antibody and TrueBlot anti-rabbit beads for 3 h at 4°C.

For FLAG immunoprecipitations, HEK293 cells were transfected with the indicated FLAG-tagged constructs, lysed after 24h, and incubated overnight at 4°C with anti-FLAG M2 magnetic beads. Beads were washed three times with TG lysis buffer and bound proteins were eluted in 2× sample buffer.

For GFP-Trap immunoprecipitation, HeLa Flp-In T-REx EGFP-DYRK1B cells were treated with tetracycline and, where indicated, DYRK inhibitors before lysis from 10-cm dishes in TG buffer. Clarified lysates were incubated with GFP-Trap magnetic beads (Chromotek) for 2h at 4°C and washed three times before elution in sample buffer.

For V5 immunoprecipitation from PaTu 8988T cells stably expressing V5-tagged DYRK1B, cells were lysed in PBS containing 1% Triton X-100 and protease inhibitors. After centrifugation (10,000 × g, 10 min, 4°C), lysates were pre-cleared with rabbit IgG and Protein A magnetic beads and then incubated for 4h at 4°C with anti-V5 antibody or control rabbit IgG together with magnetic beads. Beads were washed five times in lysis buffer and boiled in 1× Laemmli buffer.

### In vitro kinase assays and targeted phosphosite mapping

For in vitro kinase assays, HEK293 cells were transfected with pEGFP-C1-DCP1A constructs and EGFP-DCP1A proteins were isolated 24h later by immunoprecipitation.

Bead-bound EGFP-DCP1A was incubated with recombinant GST-DYRK1B (Invitrogen PV4649) in kinase buffer containing 50 mM Tris-HCl pH 7.5, 0.1 mM EGTA, 0.1% 2-mercaptoethanol, 10 mM MgCl2 and 0.1 mM ATP for 60min at 30°C, with or without AZ191 (1 μM) or harmine (1 μM). Reactions were terminated by boiling in Laemmli buffer and analysed by immunoblotting.

For targeted mapping of DCP1A phosphorylation sites, three DCP1A preparations were analysed: endogenous DCP1A immunoprecipitated from HD1B cells after tetracycline treatment (24h), EGFP-DCP1A transiently expressed in HD1B cells before tetracycline induction, and EGFP-DCP1A isolated from HEK293 cells and phosphorylated in vitro by recombinant DYRK1B. Immunocomplexes were resolved by SDS-PAGE, stained with Coomassie Blue and the DCP1A bands excised and split into two samples for downstream mass-spectrometric analysis, essentially as described previously (Webster & Oxley, 2005). For each sample, one gel portion was digested with trypsin and the other with chymotrypsin. Peptides were analysed by LC-MS/MS on the same system used for the SILAC analysis, but using a 0.075 x 150mm column with a 30min gradient of 0-40% acetonitrile 0.1% formic acid.

### Immunofluorescence, confocal imaging and super-resolution microscopy

For high-content quantification of processing bodies in HTetR, HD1B, HD1B(KD) and PANC-1 cells, cells were grown on glass coverslips (poly-L-lysine-coated coverslips for HTetR/HD1B derivatives), treated as indicated, washed briefly in PBS and fixed/permeabilised in ice-cold methanol for 7min. Coverslips were blocked in PBS containing 2% BSA and 0.02% sodium azide and incubated overnight at 4°C with anti-DCP1A and anti-DDX6 antibodies. After washing, coverslips were incubated with fluorescent secondary antibodies, mounted in Vectashield with DAPI and imaged at 40× on an InCell 6000 instrument. Images were analysed in CellProfiler. Foci containing DCP1A or DDX6 were identified, and processing bodies were defined as cytoplasmic foci positive for both markers. At least 150 cells were analysed per sample. For siRNA-based P-body analysis, PANC-1 cells were transfected with control or DYRK1B-targeting siRNAs, fixed 48 h later and processed as above.

For confocal analysis of DYRK family localisation, HeLa cells were transiently transfected with empty EGFP vector, EGFP-DYRK1A, EGFP-DYRK1B, EGFP-DYRK2 or EGFP-DYRK3 and fixed with 4% paraformaldehyde for 15 min at room temperature. Cells were permeabilised in 0.2% Triton X-100, blocked in 1% BSA/5% normal goat serum/0.02% Triton X-100, and incubated with anti-DCP1A, anti-DDX6 or anti-HA antibodies followed by Alexa Fluor-conjugated secondaries. Nuclei were counterstained with DAPI and coverslips were mounted in Dako fluorescence mounting medium. Images were acquired as maximum-intensity projections on a Zeiss LSM780 confocal microscope using a 63× oil-immersion objective. Line-scan colocalisation analyses were performed in Zeiss ZEN and plotted in GraphPad Prism. For super-resolution imaging of inducible HeLa Flp-In T-REx EGFP-DYRK1B cells, cells were grown on high-precision No. 1.5H coverslips, induced with tetracycline as indicated, fixed and stained for DCP1A and DDX6, mounted in ProLong Diamond Antifade and imaged on an N-SIM S microscope (Nikon) equipped with a 100×, 1.49 NA oil-immersion objective and an Andor iXon 897 camera. Three-dimensional SIM images were reconstructed in NIS-Elements and processed in Fiji.

For imaging of endogenous DYRK1B and DCP1B in PaTu 8988T cells, cells were seeded on glass coverslips. After 24 h, cells were washed with PBS and medium was changed to starving conditions (DMEM, 0.5% FBS, 1% Pen/Strep). Cells were either left untreated for 24 h, or were treated with 0.1 % DMSO (vehicle control), 1 µM LCTB21 (kindly provided by Perha Pharma (Lindberg *et al*, 2023)) or 1 µM AZ191 for 24h prior to fixing. Cells were then fixed in 4% formaldehyde for 10 min at room temperature, permeabilised with 0.5% Triton X-100 for 5min, blocked in PBS containing 10% FBS, and incubated overnight at 4°C with anti-DYRK1B and anti-DCP1B antibodies in PBS/10% goat serum/0.1% saponin, followed by Alexa Fluor-conjugated secondary antibodies for 2h at room temperature. Coverslips were mounted in Vectashield with DAPI.

For deconvolved widefield imaging in PaTu 8988T cells, z-stacks were acquired on a Leica DM5500 B widefield microscope and subjected to three-dimensional deconvolution and maximum-intensity projection. Quantification of DCP1B-positive PBs from these images was performed in ImageJ.

STED imaging was performed on a Leica STELLARIS STED microscope using an HC PL APO CS2 93×/1.30 GLYC objective. Excitation was provided by the white-light laser and a 405 nm laser as appropriate for the fluorophores used. Images were acquired with a theoretical pixel size of 60 nm, with depletion at 592 nm and 775 nm and photon-counting τSTED detection.

### Live Cell Imaging

For live cell imaging of DYRK1B and PBs, PaTu 8988T cells were transiently co transfected with DYRK1B-GFP and mCherry-DDX6 (generated using 25033 and 54563 plasmids from Addgene) expression constructs using polyethylenimine (PEI) (Polysciences). The culture medium was replaced after 24h with fresh DMEM supplemented with 10% FBS and 1% penicillin/streptomycin prior to imaging.

Live-cell imaging was performed on a Leica STELLARIS STED microscope (Leica Microsystems, Wetzlar, Germany). Confocal images were acquired to monitor the dynamics of DYRK1B-GFP and mCherry-DDX6. Imaging was carried out using the same fluorescence excitation and detection settings as described for super-resolution microscopy, but without application of STED depletion lasers.

### Quantifications and statistics

High content imaging data presented in this study were derived from at least three independent biological replicates. Image analysis, graph generation, and statistical tests were performed using GraphPad Prism (versions 8–10; GraphPad, San Diego, CA). Details of data analysis are provided in the corresponding Figure legends.

Processing body quantification from CellProfiler-based assays was analysed using one-way ANOVA with appropriate multiple comparison tests, as indicated in the Fig legends. Quantification of DCP1B foci in pancreatic cancer cell imaging experiments, including knockout and rescue conditions, was analysed using unpaired, two-tailed t-tests.

Data are presented as mean ± standard deviation (SD) or standard error of the mean (SEM), as specified in the Figure legends.

Phospho-SILAC data was analysed generating a volcano plot detailing the unadjusted -log_10_ (p-values) and average log_2_ (ratios) for phosphopeptides (filtered for those quantified in at least 2 experiments). Significantly differentially abundant phosphopeptides were defined as those with a fold change >1.5, p-value <0.05.

The StringDB network (version 12.0) of those proteins with significantly higher phosphopeptide abundance in treated vs untreated (interaction confidence >0.4) and StringDB used for the gene ontology annotation of proteins in this network. Gene ontology term enrichment was determined by DAVID Bioinformatics (v2023q2 Knowledgebase) for proteins with peptides exhibiting a mean 1.5-fold increase.

## SUPPLEMENTARY INFORMATION

**Supplementary Table S1 DYRK1B SILAC phosphoproteomics**

**Supplementary Table S2 DCP1A targeted phosphoproteomics**

**Movie EV1**- Live-cell confocal imaging of PaTu 8988T cells transiently expressing DYRK1B-GFP and mCherry-DDX6. The movie shows cytoplasmic DYRK1B positive puncta overlapping with DDX6 positive at PBs.

**Figure EV1 (related to Fig. 1):**
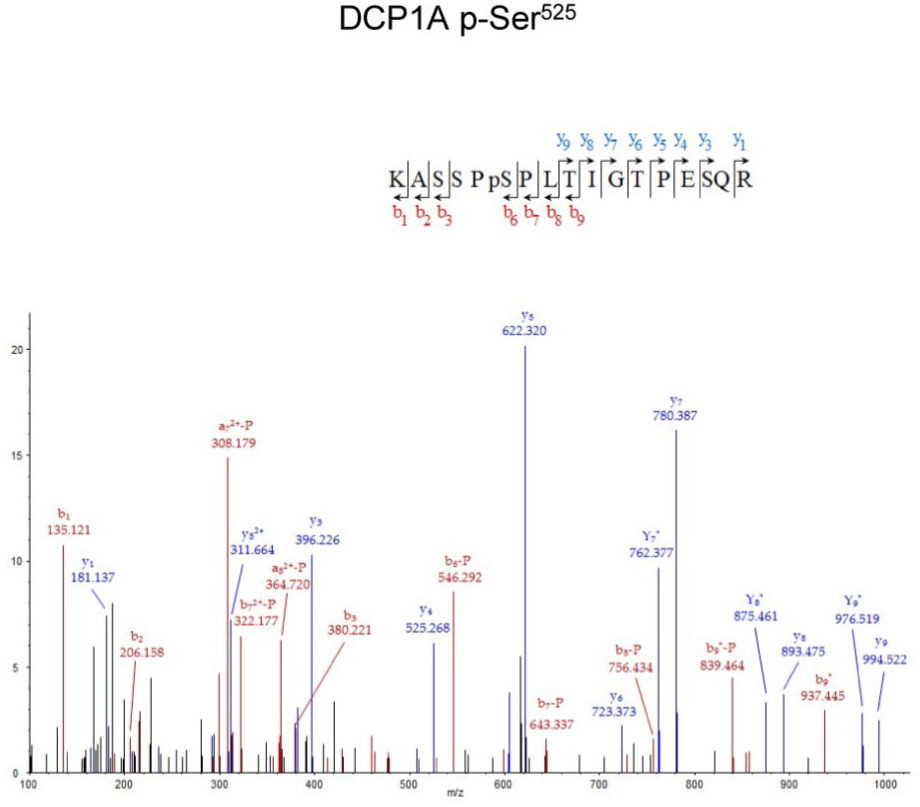
Spectrum graph from SILAC mass spectrometry data showing phosphorylated DCP1A peptides.

**Figure EV2 (related to Fig. 2):**
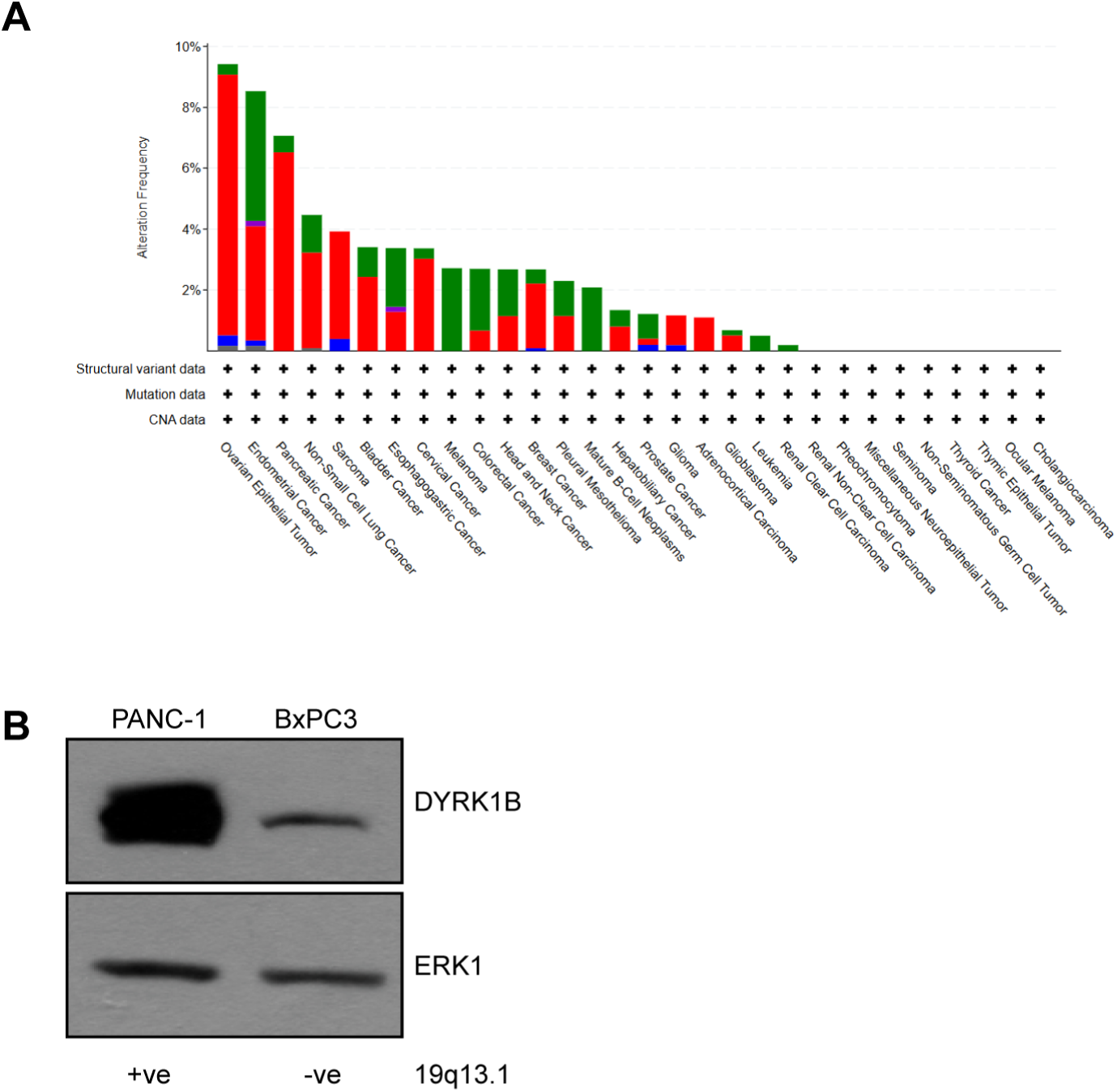
DYRK1B amplifications in cancer. (A) Graph obtained from c-bioportal, showing 7% of pancreatic cancer show DYRK1B amplifications. (B) Western blot showing pancreatic cancer cell line (PANC-1) which harbours 19q13.1 amplification exhibiting higher levels of DYRK1B compared to a different pancreatic cancer cell line BxPC3 which does not have 19q13.1 chromosomal amplification.

**Figure EV3 (related to Fig. 4):**
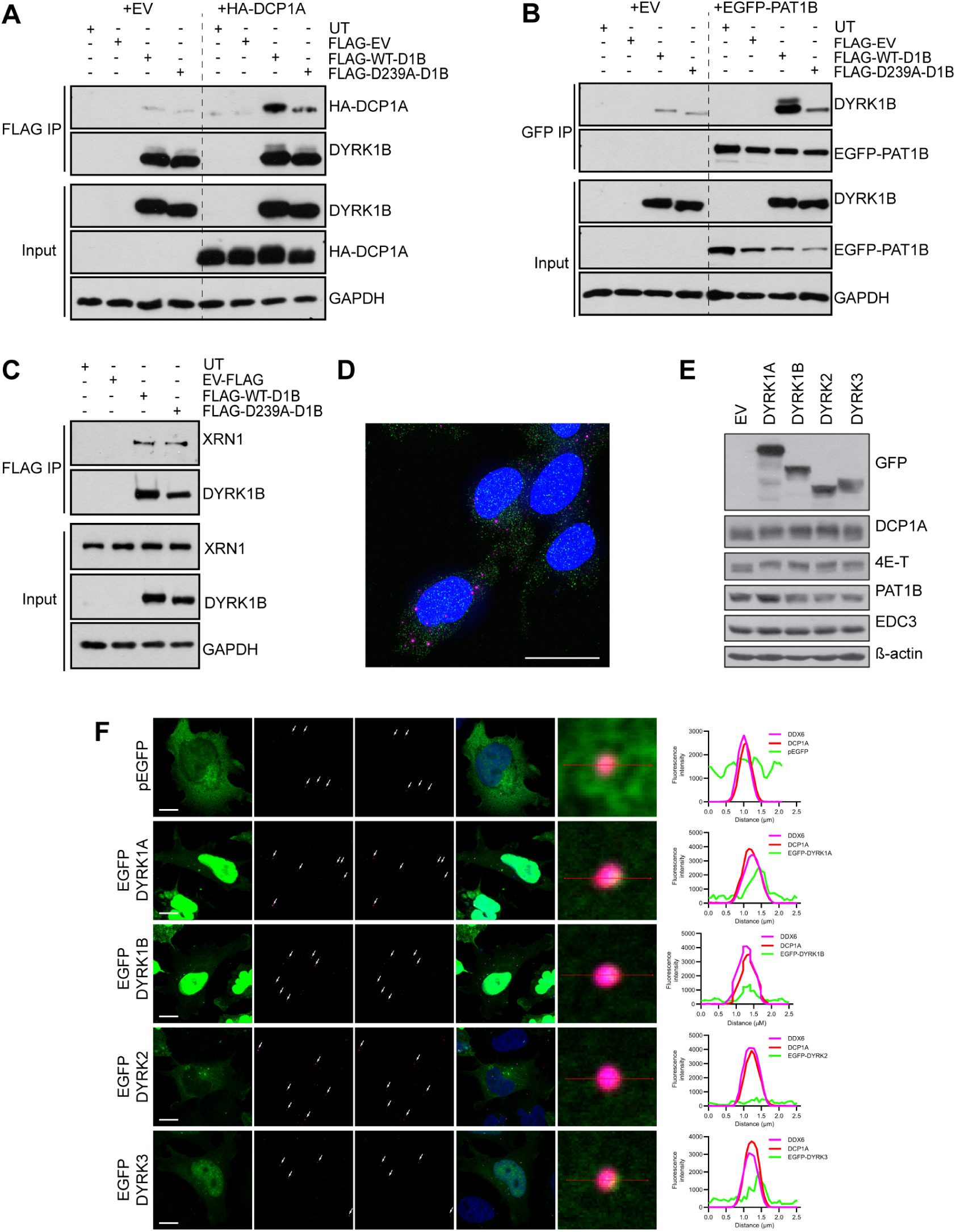
P-body components interact with and co-localise with DYRK1B in PBs. (A) FLAG IP of FLAG-WT-D1B or FLAG-D239A-D1B transfected HEK293T cells co-transfected with empty vector (EV) control or HA-DCP1A associates with HA-DCP1A. (B) GFP trap IP of FLAG-WT-D1B or FLAG-D239A-D1B transfected HEK293T cells co-transfected with empty vector (EV) control or EGFP-PAT1B associates with both WT and kinase dead DYRK1B. (C) FLAG IP of FLAG-WT-D1B or FLAG-D239A-D1B transfected HEK293T cells associates with endogenous XRN1. (D) 3D-deconvoluted widefield image of PaTu 8988T cells stained for endogenous DYRK1B (green) and DCP1B (magenta) showing colocalization. Scale bar in the whole cell image represents 20µm. (E) HEK293 cells transfected with EGFP tagged DYRK1A, DYRK1B, DYRK2, DYRK3 showing varying levels of phosphorylation of DCP1A, 4E-T, PAT1B, EDC3 proteins. (F) Confocal images of HeLa cells transiently transfected with either control empty vector EGFP, WT EGFP-DYRK1B, EGFP-DYRK1A, EGFP-DYRK2, EGFP-DYRK3 and stained for DDX6 (red) and DCP1A (magenta). Scale bars, 10µm. Line scan analyses of fluorescence intensities of PBs (DDX6 and DCP1A) along with the various EGFP-tagged DYRK vectors.

